# Loss of premature stop codon in the *Wall-Associated Kinase 91* (*OsWAK91*) gene confers sheath blight disease resistance in rice

**DOI:** 10.1101/625509

**Authors:** Noor Al-Bader, Austin Meier, Matthew Geniza, Yamid Sanabria Gongora, James Oard, Pankaj Jaiswal

## Abstract

The genetic arms race between pathogen and host plant is a tug of war that has been ongoing for millennia. The “battles” are those of disruption, restoration of signaling and information transmission on a subcellular level. One such battle occurs between rice an important crop that feeds 50% of the world population and the sheath blight disease (SB) caused by the fungus *Rhizoctonia solani*. It results in 10□30% global yield loss annually and can reach 50% under severe outbreak. Many Receptor□like kinases (RLKs) are recruited as soldiers in these battles. Wall Associated Receptor Kinases (WAKs) a subfamily of receptor-like kinases have been shown to play a role in fungal defense. Here we show that rice gene *OsWAK91*, present in the major SB resistance QTL region on Chromosome□9 is a key component in defense against rice sheath blight. An SNP mutation C/T separates susceptible variety, Cocodrie (CCDR) from the resistant line MCR010277 (MCR). The resistant allele C results in the stop codon loss that results in 68 amino acids longer C□terminus carrying longer protein kinase domain and phosphorylation sites. Our genotype and phenotype analysis of the top 20 individuals of the double haploid SB population shows a strong correlation with the SNP. The susceptible allele appears as a recent introduction found in the japonica subspecies reference genome and a majority of the tropical and temperate japonica lines sequenced by the 3000 rice genome project. Multiple US commercial varieties with japonica background carry the susceptible allele and are known for SB susceptibility. This discovery opens the possibility of introducing resistance alleles into high yielding commercial varieties to reduce yield losses incurred by the disease.

## Introduction

The genetic arms race between pathogen and host plant is a tug-of-war that has been going on for millennia. The “battles” are those of disruption, restoration of signaling and information transmission on a subcellular level. One such battle occurs between rice, an important crop that feeds 50% of the global population^1^ and the soil-borne pathogen *Rhizoctonia solani* that causes the leaf sheath blight disease (SB). Leaf sheath blight is a major disease in rice that negatively affects crop yield and quality^2^. Identified by the lesions on the leaf sheath, SB causes leaves and tillers (secondary shoots) to undergo early senescence, drying out and tissue death. Ultimately, the significant loss in leaf area due to infection and death affects the plant’s photosynthetic ability, leading to reductions in biomass and yield. There are no known rice cultivars fully resistant to leaf sheath blight disease^3–5^. In the United States, planting high-production rice varieties that are susceptible to SB, resulted in up to 50% yield losses^3^, in a severe breakout season. As a result, about 72% of the rice planting area in the United States is sprayed with fungicide that incurs unsustainable economic cost to the farmers^6^. The spraying may also pose severe ecological constraints on animals and microbiome that co-inhabit and/or frequently visit rice farms and fresh water bodies. Classified as a soilborne basidiomycete fungus, *R. solani* is a particularly destructive plant pathogen^7^. In addition to its impact on rice, *R. solani* infects other major crops such as soybean, barley, sorghum, tomato, and maize^8^. Although *R. solani* rarely produces spores for a mode of mobility, sclerotia are formed which can survive in the soil for up to two years^9^. Flooding in paddy fields, as common practice or natural occurrence combined with a highly humid environment, the sclerotia spread and attach to the plant, causing SB in the susceptible varieties.

Currently, disease management for SB relies heavily on fungicides. This approach may not be the most sustainable due to potential resistance developing in *R. solan*i populations, and pollution of agricultural resources^8^. For example, in 2012, a strain of *R. solani* was isolated in the state of Louisiana, USA which was resistant to the strobilurin class of fungicides used by rice farmers^10, 11^. A more sustainable approach to disease management is to breed genetic resistance to SB in commercial varieties. In this study, mRNA sequencing was utilized to compare gene expression profiles of resistant and susceptible rice plants inoculated with *R. solani*. Using the transcriptome profiles of two rice lines, susceptible Cocodrie (CCDR)^12^, and resistant MCR010277 (MCR)^13^ (Figure-1A), we examined the response to *R. solani* infection at multiple time points including Day-0 (untreated control), and Day-1, 3, and 5 after inoculation.

**Figure-1:**
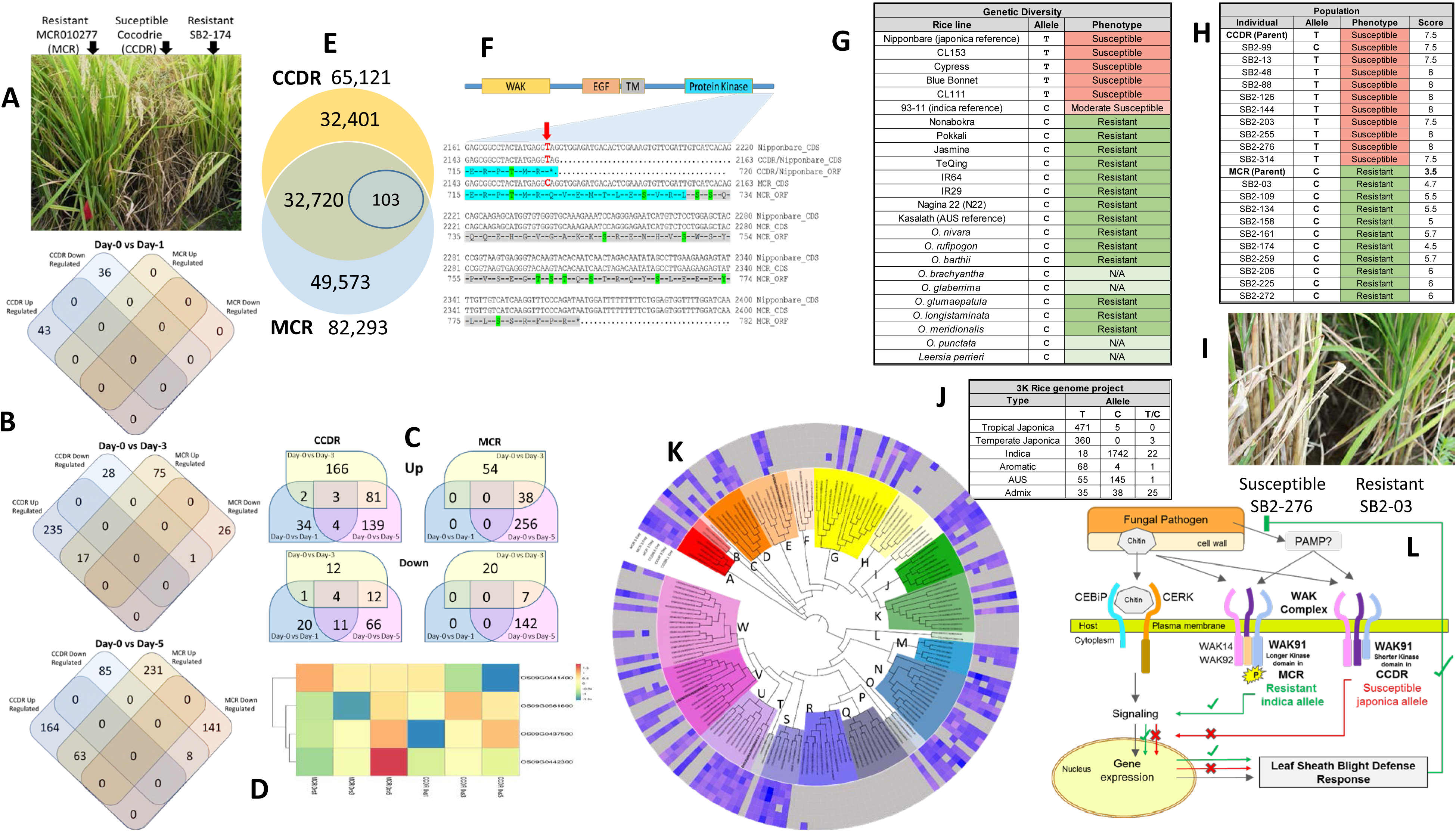
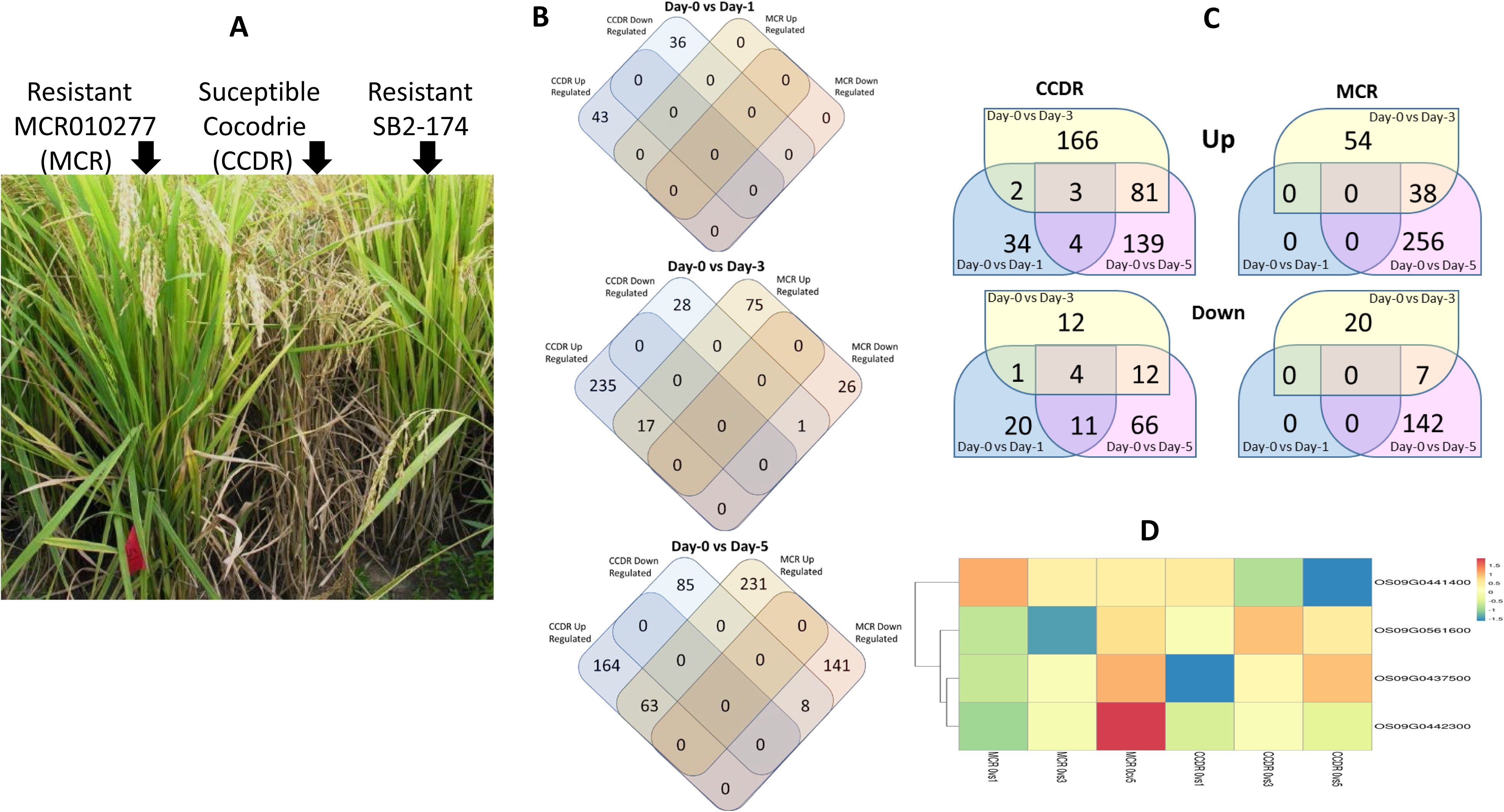
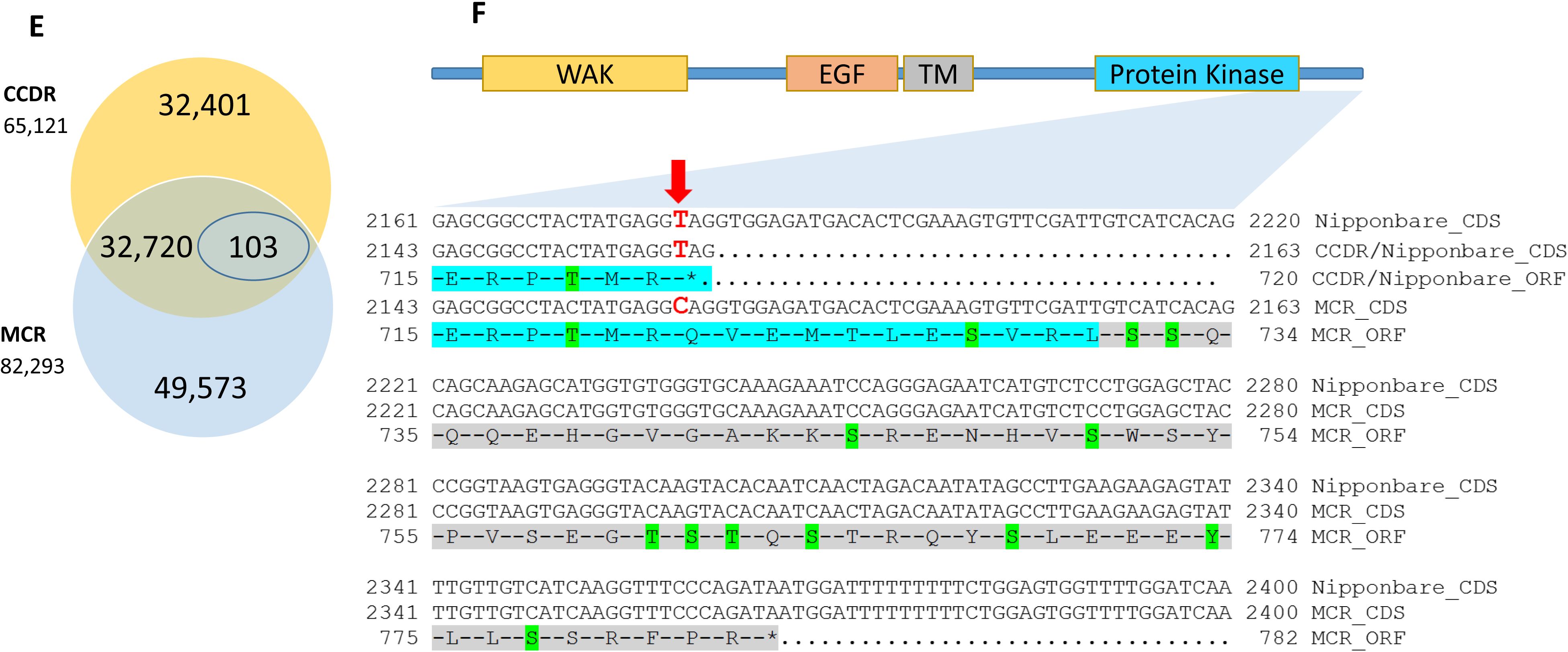
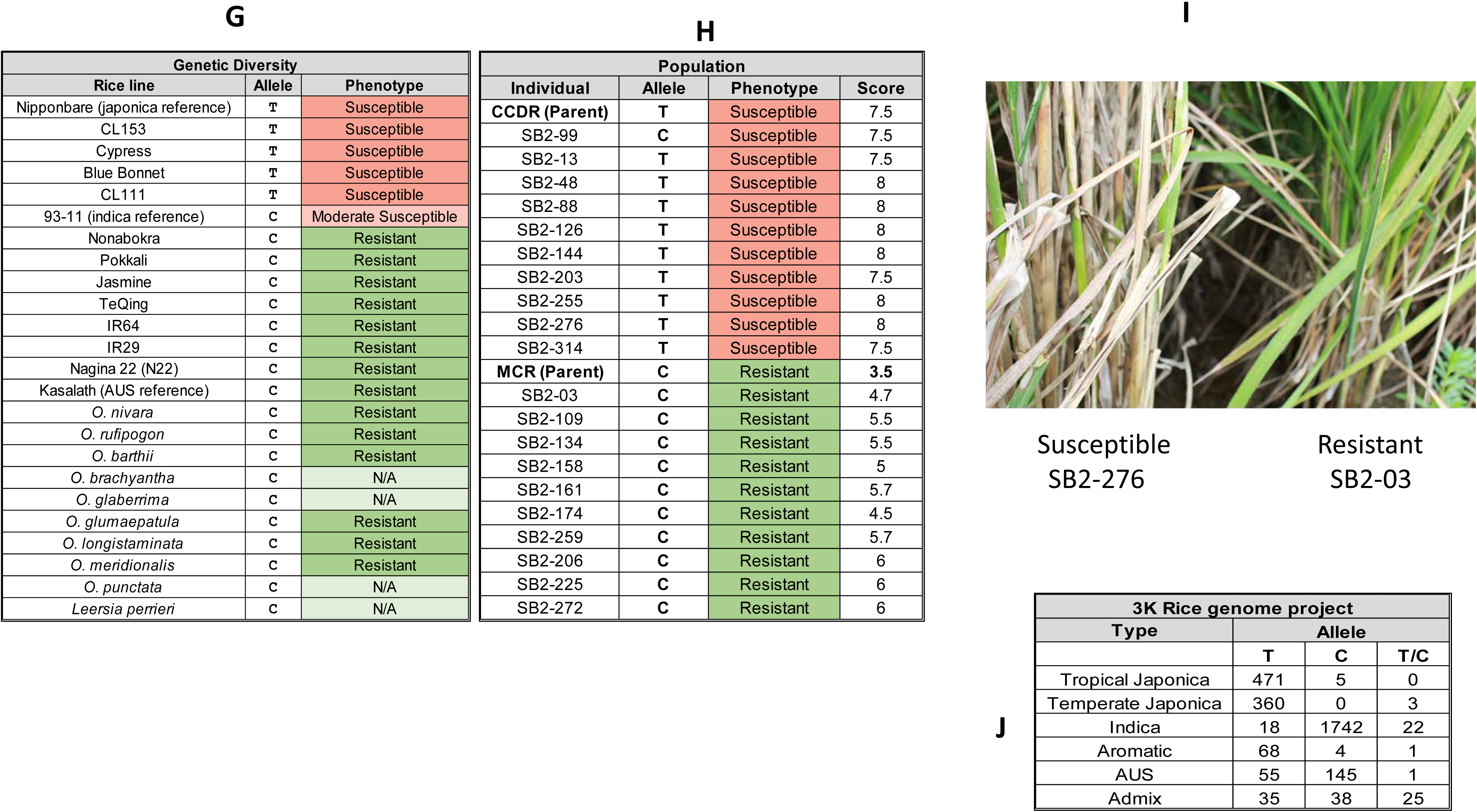
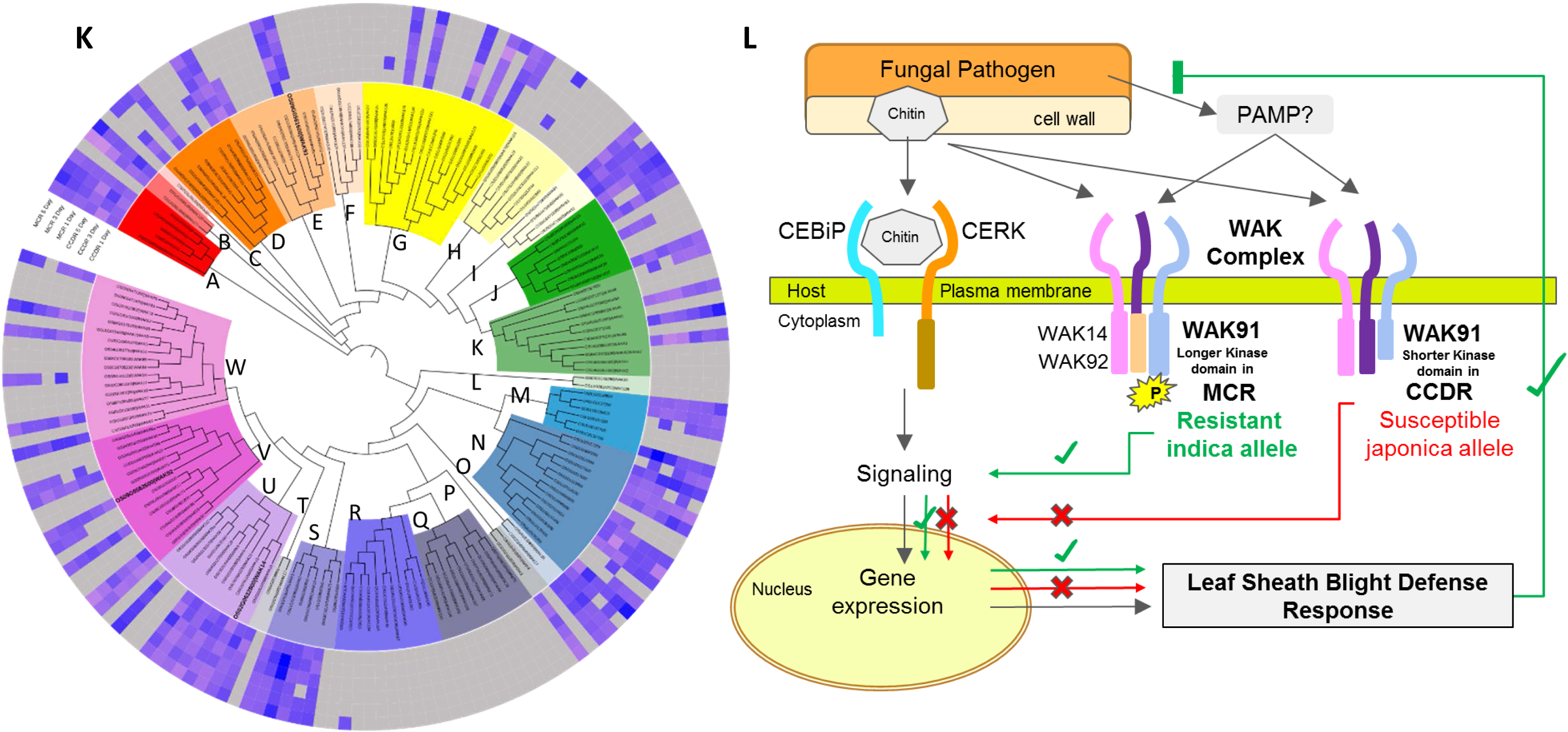
(A) The rice leaf sheath blight disease response shown by the resistant parent MCR010277 (MCR), the susceptible parent Cocodrie (CCDR) and the SB2-174 individual of the SB2 double haploid population. The disease response was tested by infecting the plants with the fungal pathogen *Rhizoctonia solani* (strain LR172). (B) The number of differentially expressed genes between the two parent lines when compared to the Day-0 vs Day-1, Day-3 and Day-5 after infection. (C) Number of up and down regulated genes shared between the time points within each rice line. (D) The differential expression of four candidate genes overlapping the major leaf sheath resistance QTL on Chromosome 9 of rice. Rows are each of the four candidate genes and columns are (L-R) first three, MCR Day-0 vs Day-1, Day-0 vs Day-3, Day-0 vs Day-5 and last three, CCDR Day-0 vs Day-1, Day-0 vs Day-3, and Day-0 vs Day-5 (E) Number of unique and shared SNPs and indels identified in the CCDR and the MCR lines. (F) The OsWAK91 ORF showing the regions of the coded domains (top) and zoom-in view of the CDS and the ORF from the reference Nipponbare, and parents CCDR and MCR rice lines (bottom). The presence of allele C in the leaf sheath blight resistant MCR line results in stop codon loss and 82 amino acid longer WAK91 peptide which carries a longer protein kinase domain(cyan colored) and additional predicted sites for serine (S), threonine (T) and tyrosine (Y) phosphorylation sites (green colored). (G) Genotyping and Phenotyping of the top ten resistant and ten susceptible individuals of the SB2 double haploid population. The population and the two parents were evaluated for sheath blight response phenotype on a scale of 0-9, where 0 = no disease and 9 = dead plant. The OsWAK91 SNP alleles T/C were scored for presence/absence. (H) OsWAK91 SNP allele T/C in 13 rice lines was confirmed by sequencing the SNP. Synteny-based sequence data was mined at the publicly available Gramene database for the reference lines indica 93-11, japonica Nipponbare, the wild species of *Oryza* genus and its outgroup *Leersia Perrieri*. The leaf sheath blight disease response phenotypes were mined from the previously published literature. (I) The leaf sheath blight response phenotype of the individuals SB2-276 (susceptible) and SB2-03 (resistant) from the DH population. (J) Occurrence of the OsWAK91 SNP allele T/C The genotyping for in the genomes of the 3000 rice lines and their grouping (K) Phylogenetic tree of the rice Wall associated Kinase gene family members distributed in 23 Subfamilies. Differential gene expression data from the MSC and CCDR rice lines were plotted in the track order (outside to inside) MCR Day-0 Vs Day-1, Day-3, Day-5 and CCDR Day-0 Vs Day-1, Day-3 and Day-5. (K) (K) A model of the host-pathogen interaction showing response to fungal chitin binding by the chitin binding receptor complex, and potential interactions and functions by the heterotrimeric WAK receptor protein complex. The WAK complex is functional in both the MCR and the CCDR rice lines, however the MCR OsWAK91 C-terminus carrying the longer protein kinase domain and the additional serine, threonine and tyrosine phosphorylation sites is expected to play a role in successfully initiating the downstream signaling response to provide leaf sheath blight resistance.

The high-quality rice reference genome of *O. sativa japonica* cv Nipponbare (IRGSP v1.0)^14^, bioinformatics analysis tools and sequences from public databases were used to profile variety and time point-specific differential expression of all rice genes with special focus on those overlapping and around the known major SB-resistance QTL on Chromosome 9^15^. We aligned the transcriptome sequence reads to the reference rice genome to identify variety-specific synonymous and non-synonymous SNPs, indels and additionally probed their putative consequences on transcript structure, regulation and protein function *in-silico*. The select set of SNP markers overlapping the differentially expressed candidate genes from the major SB-resistance QTL region on chromosome-9 were confirmed by sequencing. The selected markers were also genotyped in resistant and susceptible individuals derived from a biparental (CCDR x MCR) doubled haploid (DH) population of 197 individuals.

We observed clear gene expression and phenotype differences in response to *R. solani* infection in each of the two rice parental lines. Our findings also reveal plant defense response genes that are differentially expressed in CCDR and MCR following inoculation and carry genetic variation that alters the gene function. Based on the gene expression data, genotyping and phenotyping, we present strong evidence supporting a rice *Wall Associated Kinase 91* (Os*WAK9*1) gene as a potential breeding target for developing SB resistant breeding lines.

## Results

### Transcriptome analysis

Twenty-four cDNA libraries were created, with 3 replicates from 4 time points for each rice line, Day 0 (untreated control) and Day-1, 3, and 5 after *R. solani* inoculation. Strand-specific 150bp paired-end sequencing of the cDNA libraries yielded 308,482,684 (CCDR) and 357,018,002 (MCR) reads. After filtering the low-quality raw reads, we obtained 299,687,236 and 346,498,184 (∼98.5% of total) from the CCDR and MCR samples respectively, with an average of 27.7 million reads per sample. The cDNA sequence reads were aligned to the *O. sativa japonica* cv Nipponbare (IRGSP v1.0) reference genome. We found 16,480 (CCDR) and 17,793 (MCR) protein coding genes which show normalized expression from a minimum of one time point. Expression of known genes from each line, at each time point, was compared in a pairwise fashion against the untreated, Day-0 sample for differential gene expression analysis. For CCDR there were 105, 367 and 377 transcripts from 79, 281 and 320 genes at the Day-0 to Day-1, Day-0 to Day-3, and Day-0 to Day-5 comparisons, respectively. Whereas, in the MCR line, there were 148 and 597 transcripts from 119 and 443 genes in the Day-0 to Day-3, and Day-0 to Day-5 comparisons, while there were no statistically significant differences in expression observed at Day 0 to 1 (Figure-1B). In the Day-0 to Day-3 comparison, 17 common genes were up-regulated in both the CCDR and MCR lines, and one common gene was down-regulated. Similarly, in the Day 0 to 5 comparison, 63 common genes in the CCDR and MCR lines were up-regulated, while 8 genes were down-regulated (Figure 1B). In the susceptible CCDR line, there was a larger number of differentially-expressed genes at Day-1, and increasing numberson and Day-3, unlike the MCR line which has larger number of differentially expressed genes on Day-5 (Figure 1C).

Genes up-regulated in the CCDR line include transcription factors, and genes playing a role in the biotic stress response (both at Day-0 to 1), and enrichment of hydrolase function in the Day-0 to Day-3 comparison. One gene (OS02G0129800) which is grass family-specific, was down-regulated in both CCDR and MCR lines. This gene is known to be up-regulated in response to *Burkholderia glumae*^16^ the causal agent of rice bacterial panicle blight disease, whereas it is down-regulated in response to *Xanthomonas oryzae* the causal agent of rice bacterial leaf streak disease^17^. The Day-0 to Day-5 comparison shows enrichment of genes associated with catabolism of peptidoglycan, an important cell wall component, those associated with biotic stimulus and defense responses, kinases and those with chitinase enzyme activity required for degrading the fungal pathogen cell walls.

We found 1948 and 1914 genes from the CCDR and MCR lines, respectively, that undergo alternative splicing of transcripts. Of these, 1890 genes were common to both lines. Of all the genes showing alternative splicing under biotic stress, 57 and 20 unique genes were differentially-expressed in the CCDR and MCR lines, respectively. Intron retention was identified as the most prevalent form of transcript splicing event in response to *R. solani* infection (Table-1). This is consistent with known reports from plants, including those under abiotic stress^18^.

**Table-1.** Summary of the number of genes and events undergoing alternative splicing of transcripts in the susceptible CCDR and resistant MCR rice lines at different time points.

| Rice line | MCR |  |  |  |  |  |  |  |
| --- | --- | --- | --- | --- | --- | --- | --- | --- |
| Time points | Day-0 |  | Day-1 |  | Day-3 |  | Day-5 |  |
| Splicing type | Genes | Events | Genes | Events | Genes | Events | Genes | Events |
| Alt 5' UTR | 335 | 392 | 339 | 396 | 335 | 392 | 335 | 392 |
| Alt 3' UTR | 553 | 652 | 553 | 652 | 551 | 650 | 551 | 650 |
| Exon Skipping (ES) | 189 | 204 | 191 | 207 | 185 | 200 | 185 | 200 |
| Intron retention (IR) | 1,357 | 1,651 | 1,357 | 1,651 | 1,350 | 1,644 | 1,351 | 1645 |
| Rice line | CCDR |  |  |  |  |  |  |  |
| Time points | Day-0 |  | Day-1 |  | Day-3 |  | Day-5 |  |
| Splicing type | Genes | Events | Genes | Events | Genes | Events | Genes | Events |
| Alt 5' UTR | 337 | 394 | 336 | 393 | 335 | 392 | 336 | 393 |
| Alt 3' UTR | 557 | 656 | 553 | 652 | 551 | 650 | 553 | 652 |
| Exon Skipping (ES) | 186 | 201 | 189 | 204 | 187 | 202 | 185 | 200 |
| Intron retention (IR) | 1,371 | 1,666 | 1,369 | 1,664 | 1,355 | 1,649 | 1,357 | 1,651 |

Four differentially-expressed candidate genes were identified in the vicinity of the major SB resistance QTL on chromosome-9; a dormancy auxin-associated family protein (OS09G0437500), an elicitor-inducible cytochrome P450 (OS09G0441400), a cysteine peptidase (OS09G0442300) and a *Wall-Associated receptor kinase 91* (*WAK91;* OS09G0561600) with contrasting expression profiles (Figure 1D). The *WAK91* transcript showed down-regulation up to Day-3, followed by up-regulation at Day-5 in the MCR line compared to being constitutively expressed in the CCDR line. OS09G0441400, the elicitor-responsive P450 gene, showed an expression profile opposite that of *WAK91*. OS09G0437500, the dormancy auxin-associated gene, exhibited a similar expression profile between the two lines, except at Day-1 in CCDR when it was down-regulated. In contrast, OS09G0442300, the cysteine peptidase, was highly expressed in MCR at Day-5.

### SNP marker discovery and variant prediction

Using the transcriptome sequence reads, we identified 65,121 and 82,293 genetic variants, including single nucleotide changes (SNPs in the form of transitions and transversions and insertions and/or deletions (indels) in the susceptible CCDR and the resistant MCR lines, respectively. A total of 32,401 and 49,573 SNPs were unique to CCDR and MCR, respectively, and they shared 32,720 SNPs present at the same loci. Of the shared loci, 103 SNP sites contained alleles different from the reference and between the two lines (Figure 1E). On average, we observed that the MCR line carries ∼2500 transitions (Ts), ∼750 transversion (Tv) and ∼1000 more indel SNP events compared to the CCDR line (Table-2). In general TvSNPs are have been shown to have larger regulatory effects compared to the TsSNPs^19^, and rice is known to carry more of the transition types of SNPs^20^. The TsSNPs were greater in number in both lines, but the transition to transversion ratio (Ts/Tv) in the MCR line was 2.8 compared to 2.68 in CCDR. Within the TsSNPs, there were almost the same number of A ↔ G, and T ↔ C transitions in CCDR compared to MCR, whereas there were more G → A and C → T transitions than A → G and T → C, respectively. The transition type G → C was high in CCDR compared to MCR, where the opposite type C → G is more abundant. There were significantly fewer TsSNP C → A in MCR (Table 2).

**Table 2.** Counts of SNPs and indels identified in the susceptible CCDR and the resistant MCR rice lines, by aligning the RNA-Seq sequence reads against the reference *O. sativa japonica* cv Nipponbare reference genome (IRGSP v1.0).

| SNP | Substitution Type | CCDR | MCR |
| --- | --- | --- | --- |
| A → G | Transition | 8,703 | 11,102 |
| T → C | Transition | 8,896 | 11,228 |
| G → A | Transition | 8,735 | 11,420 |
| C → T | Transition | 8,878 | 11,622 |
| G → C | Transversion | 2,817 | 3,523 |
| C → G | Transversion | 2,730 | 3,559 |
| T → G | Transversion | 3,274 | 4,108 |
| G → T | Transversion | 3,367 | 4,158 |
| A → C | Transversion | 3,316 | 3,964 |
| C → A | Transversion | 3,339 | 4,175 |
| T → A | Transversion | 3,734 | 4,428 |
| A → T | Transversion | 3,685 | 4,397 |
| Total SNPs |  | 61,474 | 77,684 |
| Total Indels |  | 3,647 | 4,609 |
| Total variants |  | 65,121 | 82,293 |

The identified genetic variants map to 22,448 and 24,343 rice reference protein coding genes in the CCDR and MCR lines respectively, constituting 55% and 66% of the genes in the published reference genome^14^. Because we used the cDNA sequence reads for SNP calling and genetic marker development, we used the Variant Effect Predictor (VEP) tool^21^ to predict the causal effect of each SNP on the 5’ and 3’ UTRs, exons, intron and intron splicing boundaries of the transcribed gene. If the SNP was in an exon, we also computed the synonymous (sSNP) and non-synonymous (nsSNP) variations to predict the putative consequences on transcript structure, regulation, splicing, and peptide structure and function (Table-3).

Based on these VEP results, and the differential gene expression results, we developed genetic markers that co-segregate with the disease resistance phenotype in the population study. The set of 135 SNP genetic markers^22^, including those identified herein, was short-listed by selecting for chromosomal locations overlapping the major and minor SB QTL. Sheath blight QTLs were previously identified on chromosomes 1, 2, 3, 4, 5, 6, 7, 8, 9, 11 and 12 ^15, 23–29^. In the major SB QTL region, we found that the differentially expressed *OsWAK91* (OS09G0561600) gene carried a non-synonymous SNP (T → C) aligned at position 22,318,449 bp on Chromosome-9 of the Nipponbare rice reference genome (IRGSP v1.0; Figure 1F). The SB susceptible line CCDR carried the same allele T as the reference, whereas the resistant MCR line carried the C allele. The T → C transition results in the loss of a stop codon in MCR WAK91, resulting in a predicted OsWAK91 peptide with an additional 68 amino acids. Sequencing the amplified region of the WAK91 SNP marker confirmed for presence of the susceptible T allele in the parent CCDR, as well as other US elite lines with japonica background, namely, CL53, Cypress, Blue Bonnet, and CL111, whereas in the indica lines namely, IR29, IR64, Jasmine, TeQing, Pokkali, Nonabokra, 93-11, Kasalath and Nagina22 (N22) were confirmed to carry the resistant allele C (Figure-1G).

### Population study

Evaluation of the 10 most-resistant individuals of the SB2 double haploid (DH) population produced disease ratings between 4.7 to 6.0, while the resistant parent MCR showed a 3.5 rating in the same study. In contrast, the 10 most-susceptible individuals of the population scored between 7.5 to 8.0, while the susceptible parent CCDR was a 7.5 rating (Figure-1A, 1H, 1I, Supplementary Data File 1). The selective genotyping of the parent lines MCR and CCDR and 20 selected DH lines with 135 nsSNP markers and located in the major and minor SB QTL regions on chromosomes 1, 2, 3, 4, 5, 6, 8, 9, 11 and 12. These included the non-synonymous *OsWAK91* SNP (T/C) identified in this study. An ANOVA test was performed, and the best 13 ranked nsSNP markers, based on the ANOVA analysis, F values, False Discovery Rates, and adjusted R^2^ values (Table 4), mapped to the bottom of chromosome-9, within the major SB resistance QTL that was identified previously in 12 separate studies using six *indica* lines (reviewed by Zuo et al. 2014 ^30^). In the major SB QTL region on chromosome-9, we found that the *OsWAK91* (OS09G0561600) gene carried a nsSNP (T/C) at position 22,318,449 bp on Chromosome-9 of the Nipponbare rice reference genome. Based on the genotyping data we found that, except in the SB2-99 line, the *WAK91* SNP allele C contributed by the MCR parent line always co-segregated with SB resistance phenotype, whereas the allele T from the CCDR parent co-segregates with susceptible phenotype (Figure 1H, Supplementary Data File 1).

**Table-3.**
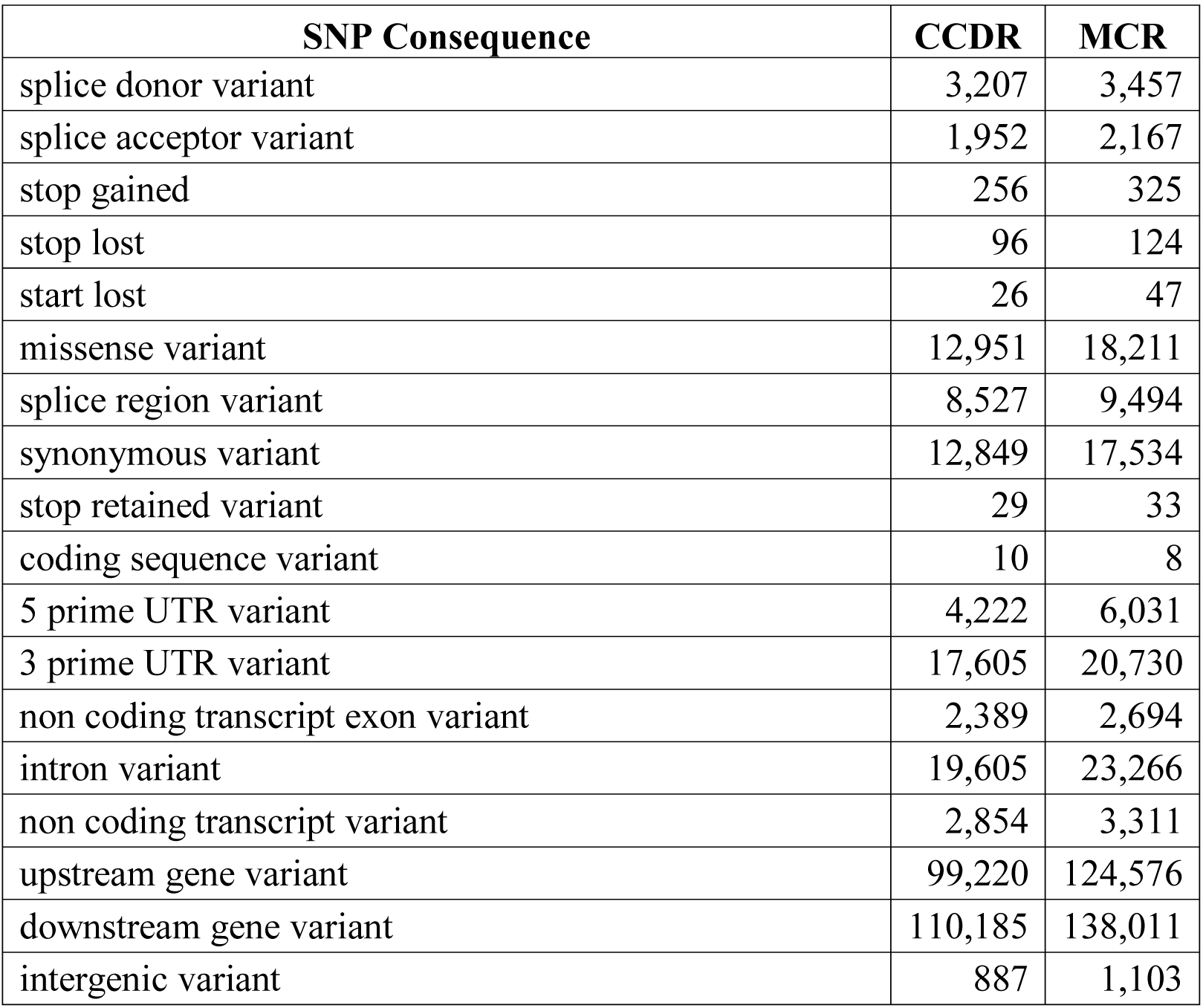
Summary of SNP consequences on gene function and structure based on its overlapping position in the genes identified in the reference *O. sativa japonica* cv Nipponbare reference genome (IRGSP v1.0).

**Table: 4.** Results of the ANOVA analysis. Includes Marker rank, MSU (v7) and IRGSP v1.0 rice gene IDs, F value, Bonferroni Hochberg False Discovery Rate (FDR), p values and R-squared values of 135 nsSNP makers genotyped in the 10 most resistant and 10 most susceptible lines of the RiceCAP project’s SB2 double haploid mapping population.

| Rank | MSU gene ID | IRGSP gene ID | F value | Raw p value | p value | FDR | R<sup>2</sup> value |
| --- | --- | --- | --- | --- | --- | --- | --- |
| 1 | LOC_Os09g32860 | N/A | 149.15400 | 0.00000 | 0.00000 | 0.00000 | 0.89232 |
| 2 | LOC_Os09g33710 | Os09g0511900 | 61.04000 | 0.00000 | 0.00004 | 0.00000 | 0.77227 |
| 3 | LOC_Os09g34180 | Os09g0517600 | 61.04000 | 0.00000 | 0.00004 | 0.00000 | 0.77227 |
| 4 | LOC_Os09g36900 | Os09g0540600 | 61.04000 | 0.00000 | 0.00004 | 0.00000 | 0.77227 |
| 5 | LOC_Os09g37230 | Os09g0544300 | 61.04000 | 0.00000 | 0.00004 | 0.00000 | 0.77227 |
| 6 | LOC_Os09g37240 | Os09g0544400 | 61.04000 | 0.00000 | 0.00004 | 0.00000 | 0.77227 |
| 7 | LOC_Os09g37590 | N/A | 61.04000 | 0.00000 | 0.00004 | 0.00000 | 0.77227 |
| 8 | LOC_Os09g37800 | Os09g0550600 | 61.04000 | 0.00000 | 0.00004 | 0.00000 | 0.77227 |
| 9 | LOC_Os09g37880 | Os09g0551400 | 61.04000 | 0.00000 | 0.00004 | 0.00000 | 0.77227 |
| 10 | LOC_Os09g38700 | Os09g0559900 | 61.04000 | 0.00000 | 0.00004 | 0.00000 | 0.77227 |
| 11 | LOC_Os09g38710 | Os09g0560000 | 61.04000 | 0.00000 | 0.00004 | 0.00000 | 0.77227 |
| 12 | LOC_Os09g38850 | Os09g0561600 | 61.04000 | 0.00000 | 0.00004 | 0.00000 | 0.77227 |
| 13 | LOC_Os09g38970 | Os09g0563250 | 61.04000 | 0.00000 | 0.00004 | 0.00000 | 0.77227 |
| 14 | LOC_Os12g10330 | Os12g0204600 | 41.63300 | 0.00000 | 0.00052 | 0.00004 | 0.69815 |
| 15 | LOC_Os12g10410 | Os12g0205500 | 41.63300 | 0.00000 | 0.00052 | 0.00004 | 0.69815 |
| 16 | LOC_Os12g13100 | Os12g0233100 | 41.63300 | 0.00000 | 0.00052 | 0.00004 | 0.69815 |
| 17 | LOC_Os12g15460 | Os12g0256900 | 41.63300 | 0.00000 | 0.00052 | 0.00004 | 0.69815 |
| 18 | LOC_Os09g39620 | Os09g0569800 | 37.14500 | 0.00001 | 0.00106 | 0.00008 | 0.67359 |
| 19 | LOC_Os12g09710 | Os12g0198900 | 33.11300 | 0.00002 | 0.00211 | 0.00016 | 0.56581 |
| 20 | LOC_Os12g10180 | N/A | 33.11300 | 0.00002 | 0.00211 | 0.00016 | 0.56581 |
| 21 | LOC_Os09g32020 | Os09g0493500 | 23.45700 | 0.00013 | 0.01449 | 0.00097 | 0.48368 |
| 22 | LOC_Os12g06980 | Os12g0167700 | 23.45700 | 0.00013 | 0.01449 | 0.00097 | 0.48359 |
| 23 | LOC_Os12g09000 | Os12g0192500 | 16.85600 | 0.00066 | 0.07303 | 0.00468 | 0.44991 |
| 24 | LOC_Os12g07950 | Os12g0179800 | 14.72200 | 0.00121 | 0.13166 | 0.00807 | 0.35200 |
| 25 | LOC_Os06g13040 | Os06g0237400 | 9.77800 | 0.00583 | 0.62916 | 0.03699 | 0.33220 |
| 26 | LOC_Os06g15170 | Os06g0262800 | 8.95400 | 0.00781 | 0.83613 | 0.04726 | 0.28421 |
| 27 | LOC_Os12g03554 | Os12g0129550 | 7.14700 | 0.01550 | 0.98350 | 0.08949 | 0.26383 |
| 28 | LOC_Os12g04660 | Os12g0140700 | 7.14700 | 0.01550 | 0.98350 | 0.08949 | 0.26383 |
| 29 | LOC_Os02g34490 | Os02g0549900 | 6.28300 | 0.02201 | 0.98350 | 0.10351 | 0.25873 |
| 30 | LOC_Os02g34850 | Os02g0554000 | 6.28300 | 0.02201 | 0.98350 | 0.10351 | 0.25873 |
| 31 | LOC_Os02g35210 | Os02g0558400 | 6.28300 | 0.02201 | 0.98350 | 0.10351 | 0.25873 |
| 32 | LOC_Os12g07800 | Os12g0177800 | 6.45100 | 0.02053 | 0.98350 | 0.10351 | 0.25598 |
| 33 | LOC_Os12g06740 | Os12g0164300 | 6.45100 | 0.02053 | 0.98350 | 0.10351 | 0.24260 |
| 34 | LOC_Os08g19694 | Os08g0293300 | 6.19300 | 0.02284 | 0.98350 | 0.10361 | 0.24120 |
| 35 | LOC_Os08g20020 | Os08g0296900 | 5.72100 | 0.02788 | 0.98350 | 0.11423 | 0.24120 |
| 36 | LOC_Os08g30850 | Os08g0399300 | 5.72100 | 0.02788 | 0.98350 | 0.11423 | 0.18705 |
| 37 | LOC_Os08g30910 | Os08g0400200 | 5.76500 | 0.02736 | 0.98350 | 0.11423 | 0.18705 |
| 38 | LOC_Os06g19110 | Os06g0295000 | 3.85200 | 0.06535 | 0.98350 | 0.20748 | 0.17626 |
| 39 | LOC_Os06g22020 | Os06g0325900 | 3.85200 | 0.06535 | 0.98350 | 0.20748 | 0.17626 |
| 40 | LOC_Os06g22460 | Os06g0330100 | 3.85200 | 0.06535 | 0.98350 | 0.20748 | 0.17626 |
| 41 | LOC_Os06g23530 | Os06g0343100 | 3.85200 | 0.06535 | 0.98350 | 0.20748 | 0.17626 |
| 42 | LOC_Os06g28124 | Os06g0475400 | 3.85200 | 0.06535 | 0.98350 | 0.20748 | 0.17626 |
| 43 | LOC_Os06g28670 | Os06g0481400 | 3.85200 | 0.06535 | 0.98350 | 0.20748 | 0.17626 |
| 44 | LOC_Os06g29700 | Os06g0492800 | 3.85200 | 0.06535 | 0.98350 | 0.20748 | 0.17626 |
| 45 | LOC_Os06g29844 | Os06g0494400 | 4.14200 | 0.05684 | 0.98350 | 0.20748 | 0.17626 |
| 46 | LOC_Os03g43684 | N/A | 4.14200 | 0.05684 | 0.98350 | 0.20748 | 0.16570 |
| 47 | LOC_Os06g31070 | Os06g0507200 | 3.57500 | 0.07486 | 0.98350 | 0.23189 | 0.12005 |
| 48 | LOC_Os06g32350 | N/A | 2.20400 | 0.15499 | 0.98350 | 0.33529 | 0.12005 |
| 49 | LOC_Os04g10460 | Os04g0183500 | 2.20400 | 0.15499 | 0.98350 | 0.33529 | 0.11751 |
| 50 | LOC_Os04g11640 | Os04g0193200 | 2.20400 | 0.15499 | 0.98350 | 0.33529 | 0.11751 |
| 51 | LOC_Os04g11970 | Os04g0196200 | 2.20400 | 0.15499 | 0.98350 | 0.33529 | 0.11751 |
| 52 | LOC_Os04g15650 | Os04g0227000 | 2.39700 | 0.13899 | 0.98350 | 0.33529 | 0.11751 |
| 53 | LOC_Os04g20680 | Os04g0275100 | 2.39700 | 0.13899 | 0.98350 | 0.33529 | 0.11751 |
| 54 | LOC_Os04g21890 | N/A | 2.39700 | 0.13899 | 0.98350 | 0.33529 | 0.11751 |
| 55 | LOC_Os04g23620 | N/A | 2.39700 | 0.13899 | 0.98350 | 0.33529 | 0.11751 |
| 56 | LOC_Os04g23890 | Os04g0304200 | 2.39700 | 0.13899 | 0.98350 | 0.33529 | 0.11751 |
| 57 | LOC_Os09g16540 | Os09g0334800 | 2.39700 | 0.13899 | 0.98350 | 0.33529 | 0.10953 |
| 58 | LOC_Os09g17600 | Os09g0344800 | 2.39700 | 0.13899 | 0.98350 | 0.33529 | 0.10953 |
| 59 | LOC_Os09g17630 | Os09g0345300 | 2.39700 | 0.13899 | 0.98350 | 0.33529 | 0.10953 |
| 60 | LOC_Os03g30130 | Os03g0415200 | 2.45600 | 0.13451 | 0.98350 | 0.33529 | 0.10907 |
| 61 | LOC_Os03g37720 | Os03g0573500 | 2.45600 | 0.13451 | 0.98350 | 0.33529 | 0.10907 |
| 62 | LOC_Os03g39150 | Os03g0588400 | 2.16600 | 0.15840 | 0.98350 | 0.33529 | 0.10907 |
| 63 | LOC_Os03g40250 | Os03g0599400 | 2.16600 | 0.15840 | 0.98350 | 0.33529 | 0.10907 |
| 64 | LOC_Os08g10560 | Os08g0206500 | 2.21400 | 0.15407 | 0.98350 | 0.33529 | 0.10739 |
| 65 | LOC_Os08g12800 | Os08g0224500 | 2.21400 | 0.15407 | 0.98350 | 0.33529 | 0.10739 |
| 66 | LOC_Os08g13870 | Os08g0236400 | 2.21400 | 0.15407 | 0.98350 | 0.33529 | 0.10739 |
| 67 | LOC_Os03g63110 | Os03g0848400 | 1.47500 | 0.24022 | 0.98350 | 0.46294 | 0.07666 |
| 68 | LOC_Os02g42412 | Os02g0636400 | 1.47300 | 0.24058 | 0.98350 | 0.46294 | 0.07575 |
| 69 | LOC_Os02g44730 | Os02g0667500 | 1.47300 | 0.24058 | 0.98350 | 0.46294 | 0.07564 |
| 70 | LOC_Os02g45160 | Os02g0673100 | 1.47300 | 0.24058 | 0.98350 | 0.46294 | 0.07564 |
| 71 | LOC_Os02g45980 | Os02g0684900 | 1.47300 | 0.24058 | 0.98350 | 0.46294 | 0.07564 |
| 72 | LOC_Os02g48210 | Os02g0712700 | 1.49400 | 0.23730 | 0.98350 | 0.46294 | 0.07564 |
| 73 | LOC_Os08g42930 | Os08g0542200 | 1.20300 | 0.28720 | 0.98350 | 0.54439 | 0.06264 |
| 74 | LOC_Os11g13650 | N/A | 1.04300 | 0.32068 | 0.98350 | 0.59891 | 0.06264 |
| 75 | LOC_Os02g58540 | Os02g0832150 | 0.92900 | 0.34792 | 0.98350 | 0.63123 | 0.05477 |
| 76 | LOC_Os01g13300 | Os01g0234100 | 0.92900 | 0.34792 | 0.98350 | 0.63123 | 0.04907 |
| 77 | LOC_Os04g56250 | Os04g0657600 | 0.89800 | 0.35586 | 0.98350 | 0.63654 | 0.04907 |
| 78 | LOC_Os08g35310 | Os08g0454500 | 0.85100 | 0.36855 | 0.98350 | 0.64118 | 0.04752 |
| 79 | LOC_Os02g39590 | Os02g0608801 | 0.85100 | 0.36855 | 0.98350 | 0.64118 | 0.04513 |
| 80 | LOC_Os02g51900 | Os02g0755500 | 0.75300 | 0.39685 | 0.98350 | 0.68108 | 0.04513 |
| 81 | LOC_Os02g52060 | Os02g0757400 | 0.61500 | 0.44303 | 0.98350 | 0.70947 | 0.04513 |
| 82 | LOC_Os02g02650 | Os02g0118700 | 0.53600 | 0.47334 | 0.98350 | 0.70947 | 0.04017 |
| 83 | LOC_Os02g43460 | Os02g0650800 | 0.53600 | 0.47334 | 0.98350 | 0.70947 | 0.03305 |
| 84 | LOC_Os02g53970 | Os02g0780200 | 0.53600 | 0.47334 | 0.98350 | 0.70947 | 0.02894 |
| 85 | LOC_Os02g54330 | N/A | 0.53600 | 0.47334 | 0.98350 | 0.70947 | 0.02894 |
| 86 | LOC_Os02g54500 | Os02g0786000 | 0.53300 | 0.47484 | 0.98350 | 0.70947 | 0.02894 |
| 87 | LOC_Os02g55180 | Os02g0795000 | 0.53300 | 0.47484 | 0.98350 | 0.70947 | 0.02894 |
| 88 | LOC_Os04g57670 | Os04g0672700 | 0.53300 | 0.47484 | 0.98350 | 0.70947 | 0.02875 |
| 89 | LOC_Os04g58720 | Os04g0683800 | 0.53300 | 0.47484 | 0.98350 | 0.70947 | 0.02875 |
| 90 | LOC_Os04g58820 | Os04g0685000 | 0.53300 | 0.47484 | 0.98350 | 0.70947 | 0.02875 |
| 91 | LOC_Os04g58910 | Os04g0685900 | 0.53300 | 0.47484 | 0.98350 | 0.70947 | 0.02875 |
| 92 | LOC_Os04g59060 | Os04g0687300 | 0.50800 | 0.48531 | 0.98350 | 0.71668 | 0.02875 |
| 93 | LOC_Os04g59540 | Os04g0691900 | 0.43300 | 0.51884 | 0.98350 | 0.74688 | 0.02875 |
| 94 | LOC_Os05g50660 | Os05g0583500 | 0.42400 | 0.52340 | 0.98350 | 0.74688 | 0.02875 |
| 95 | LOC_Os09g26300 | Os09g0432900 | 0.42400 | 0.52340 | 0.98350 | 0.74688 | 0.02743 |
| 96 | LOC_Os04g05030 | Os04g0136900 | 0.34300 | 0.56525 | 0.98350 | 0.74778 | 0.02349 |
| 97 | LOC_Os05g37040 | Os05g0442100 | 0.34300 | 0.56525 | 0.98350 | 0.74778 | 0.02299 |
| 98 | LOC_Os05g39760 | N/A | 0.35600 | 0.55839 | 0.98350 | 0.74778 | 0.02299 |
| 99 | LOC_Os08g36320 | Os08g0465800 | 0.36200 | 0.55499 | 0.98350 | 0.74778 | 0.02063 |
| 100 | LOC_Os08g36760 | Os08g0471800 | 0.37900 | 0.54578 | 0.98350 | 0.74778 | 0.02063 |
| 101 | LOC_Os09g27570 | Os09g0448100 | 0.37900 | 0.54578 | 0.98350 | 0.74778 | 0.02063 |
| 102 | LOC_Os06g44820 | Os06g0658600 | 0.37900 | 0.54578 | 0.98350 | 0.74778 | 0.01971 |
| 103 | LOC_Os02g49986 | Os02g0732600 | 0.13700 | 0.71588 | 0.98350 | 0.85611 | 0.01937 |
| 104 | LOC_Os02g09820 | Os02g0191500 | 0.13700 | 0.71588 | 0.98350 | 0.85611 | 0.01871 |
| 105 | LOC_Os02g10120 | Os02g0194700 | 0.13700 | 0.71588 | 0.98350 | 0.85611 | 0.01871 |
| 106 | LOC_Os02g56380 | Os02g0807900 | 0.12700 | 0.72610 | 0.98350 | 0.85611 | 0.00789 |
| 107 | LOC_Os02g56480 | Os02g0809100 | 0.14300 | 0.70964 | 0.98350 | 0.85611 | 0.00789 |
| 108 | LOC_Os02g57960 | Os02g0826000 | 0.14300 | 0.70964 | 0.98350 | 0.85611 | 0.00789 |
| 109 | LOC_Os06g35850 | Os06g0551800 | 0.14300 | 0.70964 | 0.98350 | 0.85611 | 0.00789 |
| 110 | LOC_Os06g37500 | Os06g0572300 | 0.14300 | 0.70964 | 0.98350 | 0.85611 | 0.00789 |
| 111 | LOC_Os01g52330 | Os01g0721300 | 0.11200 | 0.74152 | 0.98350 | 0.85611 | 0.00754 |
| 112 | LOC_Os01g52880 | Os01g0729400 | 0.11200 | 0.74152 | 0.98350 | 0.85611 | 0.00754 |
| 113 | LOC_Os01g53420 | Os01g0735900 | 0.11200 | 0.74152 | 0.98350 | 0.85611 | 0.00754 |
| 114 | LOC_Os09g25620 | Os09g0424300 | 0.14300 | 0.70964 | 0.98350 | 0.85611 | 0.00731 |
| 115 | LOC_Os09g25890 | Os09g0427800 | 0.14300 | 0.70964 | 0.98350 | 0.85611 | 0.00731 |
| 116 | LOC_Os02g10900 | Os02g0203500 | 0.13900 | 0.71698 | 0.98350 | 0.85611 | 0.00699 |
| 117 | LOC_Os05g40790 | Os05g0486300 | 0.13200 | 0.72010 | 0.98350 | 0.85611 | 0.00620 |
| 118 | LOC_Os05g41130 | Os05g0490300 | 0.13200 | 0.72010 | 0.98350 | 0.85611 | 0.00620 |
| 119 | LOC_Os05g41290 | Os05g0492200 | 0.13200 | 0.72010 | 0.98350 | 0.85611 | 0.00620 |
| 120 | LOC_Os11g19700 | Os11g0302500 | 0.04200 | 0.83928 | 0.98350 | 0.89571 | 0.00263 |
| 121 | LOC_Os11g24060 | Os11g0427700 | 0.04200 | 0.83928 | 0.98350 | 0.89571 | 0.00263 |
| 122 | LOC_Os11g24180 | Os11g0430000 | 0.04200 | 0.83928 | 0.98350 | 0.89571 | 0.00263 |
| 123 | LOC_Os11g24770 | N/A | 0.04200 | 0.83928 | 0.98350 | 0.89571 | 0.00263 |
| 124 | LOC_Os11g28950 | Os11g0479100 | 0.04700 | 0.83001 | 0.98350 | 0.89571 | 0.00263 |
| 125 | LOC_Os03g53220 | Os03g0743800 | 0.04700 | 0.83001 | 0.98350 | 0.89571 | 0.00235 |
| 126 | LOC_Os03g56400 | Os03g0775400 | 0.04700 | 0.83001 | 0.98350 | 0.89571 | 0.00235 |
| 127 | LOC_Os03g57160 | Os03g0785300 | 0.04700 | 0.83001 | 0.98350 | 0.89571 | 0.00235 |
| 128 | LOC_Os03g58390 | Os03g0798200 | 0.04700 | 0.83001 | 0.98350 | 0.89571 | 0.00235 |
| 129 | LOC_Os04g55760 | Os04g0651500 | 0.03100 | 0.86218 | 0.98350 | 0.91248 | 0.00172 |
| 130 | LOC_Os02g11820 | Os02g0208900 | 0.00000 | 0.98350 | 0.98350 | 0.98350 | 0.00007 |
| 131 | LOC_Os01g54350 | Os01g0747400 | 0.00000 | 0.98350 | 0.98350 | 0.98350 | 0.00002 |
| 132 | LOC_Os01g54515 | Os01g0748950 | 0.00000 | 0.98350 | 0.98350 | 0.98350 | 0.00002 |
| 133 | LOC_Os01g55050 | Os01g0754500 | 0.00000 | 0.98350 | 0.98350 | 0.98350 | 0.00002 |
| 134 | LOC_Os01g56040 | Os01g0765900 | 0.00000 | 0.98350 | 0.98350 | 0.98350 | 0.00002 |
| 135 | LOC_Os01g57230 | Os01g0780600 | 0.00000 | 0.98350 | 0.98350 | 0.98350 | 0.00002 |
| 136 | LOC_Os01g57900 | Os01g0788900 | 0.00100 | 0.97305 | 0.98350 | 0.98350 | 0.00002 |

## Discussion

In our quest of finding candidate genes in rice that can provide good measure of resistance to *Rhizoctomia solani* and its application for breeding improved disease resistant rice, our study integrated transcriptomics, identifying genetic variation and its causes on the gene function. We identified the rice candidate gene *Wall-Associated Kinase 91* (*OsWAK91*) carrying a T/C mutation at the stop codon. The *OsWAK91* SNP site at position 22,318,449 bp on Chromosome-9 of the reference rice genome is part of stop codon (TAG) common to CCDR and the reference japonica line Nipponbare. Due to the presence of allele C in the resistant line MCR, the stop codon TAG is lost and replaced by the codon CAG in the OsWAK91 open reading frame. The codon CAG codes for glutamine, a polar uncharged side chain amino acid. The stop codon loss in the MCR line also results in a OsWAK91 peptide that is 68 amino acids longer compared to that of CCDR and Nipponbare (Figure 1F). These results lead to two hypotheses: (1) The *OsWAK91* SNP allele C is associated with SB resistance that is evolutionarily conserved with *Oryza* ancestors, and (2) the longer OsWAK91 ORF resulting from the loss of the stop codon results in a peptide with gain in novel molecular function which may play a role in leaf sheath blight resistance.

To test if the association of the *WAK91* SNP allele C with SB resistance is evolutionarily conserved, we sequenced a PCR-amplified fragment overlapping the *WAK91* SNP loci from 13 additional *O. sativa* lines; five from subspecies *japonica* and eight from subspecies *indica* (Figure-1G).

All US elite rice lines have the japonica background, and are known to be SB susceptible^22^. We predicted, and confirmed that the US japonica lines CCDR, CL53, Cypress, Blue Bonnet, CL111 carry the Os*WAK91* T allele associated with known susceptibility ^4, 31, 32–38^. In contrast, the indica lines (IR29, IR64, Kasalath, Jasmin, TeQing, Nagina 22, Pokkali and Nonabokra) were confirmed to carry the resistant C allele (Figure-1G), and exhibit some level of known SB resistance phenotype ^35, 39–41^. One exception is 93-11, from which the indica reference genome was sequenced^42^, and which is known to be moderately susceptible^39, 43^. These results are consistent with the genotype and phenotype observations of the 20 individuals of the SB2 DH population (with the exception of SB2-99), in which the *OsWAK91* SNP allele C contributed by the MCR parent always co-segregated with SB resistance phenotype (Figure 1H, 1I, Supplementary Data File 1).

To gain insights on the origin of the *OsWAK91* SNP in the *Oryza* genus, we used synteny data provided by the Gramene database^44^ to query the *OsWAK91* SNP allele T/C by aligning to the sequenced genomes of the wild species; *O. longistaminata, O. glaberrima, O. punctata, O. meridionalis, O. barthii, O. glumaepatula, O. rufipogon, O. brachyantha* and the outgroup *Leersia perrieri.* All of these species carry the indica-type allele C, and the majority are known to bear the SB resistance phenotype (Figure-1G) ^3, 33, 45, 46^. This suggests that the indica-type resistant C allele is of ancestral origin. Conversely, the japonica-type susceptible T allele is a more recent introduction in the *Oryza* clade, and is restricted to the *O. sativa subspecies japonica* for its origin. We further confirmed its origin by mining the publicly-available genetic variation data from the 3000 Rice Genome Project^47^. The majority of the temperate and tropical japonica and aromatic rice lines carry the japonica T allele, whereas the majority of indica and AUS lines carry the C allele. Only a handful of the lines carry the heterozygous T/C allele (Figure 1J). We did not have access to the SB resistance phenotype data for these lines, however, based on our findings, we predict that the indica lines carrying the C allele may bear some degree of SB resistance.

The second hypothesis suggests that the *OsWAK91* gene from the SB resistant MCR parent encodes a peptide predicted to be 68 amino acid longer due to stop loss in the ORF, and this peptide may have gained a function that plays a role in providing the SB resistance phenotype. The *OsWAK91* gene is a member of the Wall-Associated receptor Kinase (WAK) gene family (Figure 1K) and codes for a plasma membrane protein. The OsWAK91 protein contains a wall-associated receptor kinase galacturonan-binding domain (WAK domain), followed by a calcium-binding epidermal growth factor (EGF)-like domain in the N-terminal half of the protein that is present on the extra-cellular side, in the middle is a single transmembrane domain spanning the plasma membrane, followed by the cytosolic side containing a protein kinase domain and additional phosphorylation sites for serine, threonine and tyrosine residues, towards its C-terminal (Figure 1F). The WAK domain is known to be linked to the pectin fraction of the plant cell wall ^48–50^.

Members of the WAK gene family are known to express and function in response to biotic and abiotic stress conditions. Expression of *Arabidopsis thaliana AtWAK1* is induced by a pathogen response, by exogenous salicylate or by its analog 2,2-dichloroisonicotinic acid, and requires the positive regulator NPR1/NIM1. The expression of complete AtWAK1 or just its kinase domain alone, can provide resistance to pathogen *Pseudomonas syringae*^51^. Maize ZmWAK-RLK a plasma membrane protein, is known for conferring resistance to northern corn leaf blight disease caused by the fungal pathogen *Setosphaeria turcica* (anamorph *Exserohilum turcicum*; previously known as *Helminthosporium turcicum*), by restricting pathogen entry into the host cells^25^. Similarly wheat WAK gene Stb6 confers fungal pathogen resistance without a hypersensitive response^52^. Barley, *HvWAK1* is known to play a role in root development; however, when compared to other cereals and Arabidopsis sequence divergence in the extracellular domain further verifies the multifunctionality of WAK genes^53^. The WAKs are known to play role in plant cell expansion during seedling development, MAP kinase signaling, and to bind to pectin polymers in the plant cell wall and have higher affinity to bind smaller pectin fibers in response to pathogen attack ^48^.

In the reference rice genome, the WAK gene family members are distributed in 23 subfamilies (Figure 1K) and are reported to play roles in development ^54^, abiotic responses ^55, 56^ and fungal disease responses ^57^. Genes from five WAK subfamilies C, F, L, and Q did not show expression in our study (Figure 1K). The gene *OsWAK1* (OS01G0136400) is known to provide resistance to the rice blast disease pathogen *Magnaporthe grisea,* and its expression is induced by salicylic acid and methyl jasmonate. OsWAK1 phosphorylates itself and OsRFP1, a putative transcription regulator which binds to the OsWAK1 kinase domain. The rice *OsWAK91* gene, also known as the *DEFECT IN EARLY EMBRYO SAC1* (*OsDEES1*), and *OsWAK1* are known for playing role in rice sexual reproduction by regulating the development of the female gametophyte (the embryo sac), and in anther dehiscence, respectively^54, 58^. Overexpression of *OsWAK25* (OS03G0225700) is known to increase susceptibility to *R. solani* and *Cochliobolus miyabeanus*, and resistance to *Xanthomonas oryzae* pv. *oryzae* (*Xoo*) and *Magnaporthe oryzae*^59^, whereas its loss of function compromises Xa21-mediated resistance ^60, 61^*. OsWAK11* has also been shown to regulates copper detoxification ^55, 56^.

Increased rice blast resistance and reduced fungal hyphae growth was observed when *OsWAK91* was overexpressed, in contrast to the overexpression of *OsWAK112* (OS10G0180800), which suppressed resistance. The genes *OsWAK14* (OS02G0632800), *OsWAK91* and *OsWAK92* (OS09G0562600) are part of the heterotrimeric WAK protein complex located on the plasma membrane in rice cells ^62, 63^ (Figure 1L). A mutant screening by Delteil et al.^57^ reported that the loss of function of *OsWAK14*, *OsWAK91* and *OsWAK92* resulted in reduced basal resistance, though it did not affect the growth and fertility^57^.. Conversely, we concluded that both the CCDR and the Nipponbare rice reference lines carry the susceptible T allele with shorter OsWAK91 ORF, and are known for SB susceptibility. We hypothesize that if the OsWAK91 function is knocked out or when the longer C-terminal is not present, the line should become increasingly pathogen susceptible. Therefore, we can expect that the Nipponbare OsWAK91 knock out line should be highly susceptible. The results published by Deltiel et al.^57^ support this hypothesis, through testing with the pathogen *Magrnaporthe grisea*, causal agent of rice blight. In their study, Tos17-transposon knockout mutant plants were generated in the japonica reference Nipponbare rice background (susceptible). The wildtype Nipponbare and the Tos-17 mutant plants always showed disease lesions in response to the pathogen, and, as expected, the *OsWAK91* mutant plant showed 2.5 fold increase in lesions.

The ability to detect and fight pathogenic microbes by the host plants is a complex process. The pattern-recognition receptors (PRRs) localized on the surface of plant cells are well-known gene family members of the Receptor-Like Kinase (RLK) proteins that assist in the recognition of PAMP (Pathogen Associated Molecular Patterns) conserved motifs from pathogens and DAMPs (Damage Associated Molecular Patterns) derived from the damage by the microbes development^64,65,63^. Also, in the case of *Magnaporthe grisea* blast disease response in rice, the chitin elicitor binding protein, OsCEBiP and a chitin elicitor receptor kinase, OsCERK, are known to form hetero-dimer complex. Upon chitin binding the CEBiP-CERK receptor complex may in turn induce transphosphorylation followed by activating downstream signaling^66^ (Figure 1L). The majority of the OsCEBiP and OsCERK gene family members show positive upregulation in the resistant MCR line (Supplementary file 1). Similarly, the OsWAKs show early transcriptional regulation induced by chitin that is partially under the control of the chitin receptor CEBiP^57^. Query of the publicly available SNP datasets ^44, 47, 67^ suggests that *OsWAK91* is known to carry ∼400 unique SNP and indels present in the 5’ and 3’ UTRs, introns, exons, and splice junctions. This includes the stop lost at position 22,318,449 bp identified by us, and indel at 9:22,315,774-22,315,791 bp leading to start lost and 5’UTR variant, and stops gained at positions 22,316,748 bp, 22,317,724 bp, 22,318,006 bp on chromosome 9 (Supplementary Data File-2). These genetic differences did not show up in our transcriptome-based SNP dataset and are potential genetic markers for further study. Therefore, based upon the supporting evidence from the genotyping, phenotyping, evolutionary ancestry of SB tolerance and the favorably segregating SNP in the SB2 DH population, we propose a working model (Figure1L) that the candidate gene *OsWAK91* provides broad tolerance to leaf sheath blight similar to when infected with pathogens *R. solani* and *Magnaporthe grisea*. The longer ORF of the *OsWAK91*in the indica lines, or the MCR genotype with allele C, OsWAK91 provides a complete kinase domain and additional active sites for serine, threonine and tyrosine phosphorylation, which are missing in the susceptible CCDR genotype, or japonica lines with shorter C-terminus (Figure 1F). OsWAK91 forms a heterotrimeric protein complex with OsWAK14 and OsWAK92 proteins, and is an essential member of the WAK protein complex. The shorter kinase domain, and C-terminus is expected to maintain proper embryo sac development and fertility in both the japonica and indica lines. The OsWAK91 in the japonica background with truncated C-terminus kinase domain is not completely dysfunctional, since, it has been shown to be involved in embryo sac development and the japonica line Nipponbare and others mentioned in this work are all fertile. Whereas, the presence of the ancestral allele C contributes to the stop lost in the candidate gene *OsWAK91*, the longer translated protein kinase domain and additional phosphorylation sites in the C-terminus in indica lines are necessary for their role in actively regulating the WAK protein complex function, and downstream signaling to affect the disease tolerance response.

## Conclusions

We determined and confirmed strong association of the *OsWAK91* SNP marker to the leaf sheath blight resistance phenotype by extending the genotyping and phenotyping study to include 10 best resistant and 10 best susceptible individuals from the double haploid population derived from the CCDR and MCR parents. We also minded the publicly available sequenced genomes of reference and ancestral *Oryza* species and the 3000 rice genome project to confirm the evolutionary source of the mutant SNP. The resistant allele and the trait appears ancestral, whereas the susceptible allele is a more recent acquisition in the *O. sativa japonica* clade. In fact all the US elite rice varieties have japonica genome integration and are known to carry the susceptibility trait. A few of them tested by us confirm the presence of the susceptible SNP allele same as the CCDR line. Our results and inferences, supported by sequencing, genotyping and phenotyping experiment are well complemented by mutant/knockout screening by earlier studies. *OsWAK91* knockout mutants make rice disease susceptible. Therefore *OsWAK91* resistant allele and the genetic marker identified by us is of ancestral origin and is a candidate for integration in the existing and new cultivated rice varieties for leaf sheath blight resistance, which may help improve global rice production by rescuing the crop affected by the pathogen *Rhizoctomia solani*

## Methods

### Plant material and Inoculation

The sheath blight resistant line MCR010277 and the susceptible variety Cocodrie (CCDR) were planted in pots placed in a mist chamber in a greenhouse located on the Louisiana State University (LSU) campus in Baton Rouge, LA. Plants were inoculated 50 days after germination with a Potato Dextrose Agar PDA medium disc (0.8 cm diameter) containing *Rhizoctonia solani* (strain LR172) mycelia. Discs were placed at the base of the stem and between the leaf blade and leaf sheath on the main culm of each plant.

Temperatures inside the mist chamber ranged from a minimum of 27_⁰_C at night to a maximum of 37_⁰_C in the day. Natural daylight was used, with day length of approximately 11 hours 30 minutes. Humidity was maintained 80-90% using a cool mist humidifier of 1.2 gallons capacity (Vicks Inc.) that was programmed to function for a two-hour period every six hours. The chamber frame was constructed with ¾ inch PVC pipe (Charlotte Pipe ^®^) covered by extra light plastic (0.31 mm) (Painter’s Plastic – Poly America). The dimensions of the chamber were: 1.32 m wide by 2.70 m length, and 1.42 m in height, (Figure 4.2) for a total capacity of 48 pots per chamber, each pot containing three plants.

Leaf samples approximately 2 cm in length were collected from the control untreated (day 0) and from the inoculation sites 1, 3 and 5 days after treatment, and placed immediately in liquid nitrogen until transfer to −80° freezer for storage.

Leaf samples from additional 13 rice lines: CL153, Cypress, Blue Bonnet, CL111 (all japonica), and 93-11, IR29, IR64, Kasalath, Jasmin, TeQing, Nagina 22 (N22), Pokkali and Nonabokra (all indica) were collected for DNA extraction, and SNP marker genotyping by DNA amplification and sequencing.

### RNA extraction and sequencing

Frozen leaf samples from the CCDR and MCR lines collected at LSU were shipped to Oregon State University on dry ice and stored at −80° - until further processing. Poly(A)-enriched mRNA libraries was prepared from total RNA extracted from the leaf samples using RNA Plant Reagent^®^ (Invitrogen Inc., USA), RNeasy kits (Qiagen Inc., USA) and RNase-free DNase (Life Technologies Inc., USA). The concentration and quality of the poly(A)-enriched mRNA was determined using a ND-100 spectrophotometer (Thermo Fisher Scientific Inc., USA) and Bioanalyzer 2100 (Agilent Technologies Inc., USA), respectively. TruSeq RNA Sample Preparation kits (Illumina Inc., USA) were used to construct sequencing libraries. An Illumina HiSeq 3000 (Illumina Inc., USA) at the Center for Genome Research and Biocomputing, Oregon State University (CGRB, OSU), was used to sequence the 150 bp Paired End cDNA libraries.

### Sequence quality control and read alignments

The program Sickle v1.33 was used to filter all reads based on read quality. Reads under the phred score 30, and read length under 150bp were rejected. FASTQ files containing quality-trimmed and filtered reads were generated, yielding MCR and CCDR high-quality reads. Sequence reads were further filtered by removing any reads showing alignment to the pathogen *Rhizoctonia solani* AG1-IB isolate (GCA_000832345.1). Quality-filtered reads were aligned to the reference *Oryza sativa* japonica cv Nipponbare (IRGSP v1.0) genome and annotation. Reads from each biological replicate, were aligned with alignment software program STAR v.2.4.1a ^68^. The resulting sequence alignment files were converted to binary alignment files (BAM), sorted coordinately and indexed using SAMTools v.13 ^69^.

### Differential gene expression (DGE)

The program DESeq2 (v 1.22.2) was used to investigate gene expression levels of the two rice lines over the time course^70^. A Python package, HTSeq v.0.6.1p1 ^71^, was used to index the alignment BAM files and generate raw counts, with a minimum quality-score cut-off of 10. The DESeq2 R-package was the used to identify differentially expressed genes. First, the raw count data was transformed using variance stabilizing transformation (vst) method of DESeq2 to filter outliers. The DESeq2 functions namely, estimate size factors and estimate dispersions, were used to normalize the aligned read counts. This was followed by the DESeq2 function to fit a negative binomial general linear model and Wald test statistics to detect the significance scores (p-value). The final set of differentially expressed transcripts were called at the FDR, Benjamin Hoffman, corrected cutoff P-value of 0.05. Furthermore, the non-significant events identified by the DESeq2 software were filtered out. Principle components (PCA) plots were used to visualize cross-sample comparison in each DE pairwise comparison. MA-plots were generated to visualize the different between signal intensity between samples in each comparison and dispersion plots were generated to visualize the fitted normalized counts.

### SNP discovery and variant prediction

Indexed BAM files were analyzed to identify genetic variations including single nucleotide changes (SNPs; transitions and transversions) and insertions and/or deletions (indels). Alignments from each variety were pooled using the mpileup function of SAMtools^69^. VarScan.v2.3.9 (release 80)^72^ was used to identify SNPs and indels with four different minimum read coverage of categories of 2, 6, 8, and 20 and default minimum variant frequency of 0.8 with a p-value of 0.005. A consensus set of SNPs were identified in all four minimum read coverages. A genotype-specific SNP output data file in VCF format was generated. The Variant Effect Predictor (VEP) workflow provided by the Gramene database^21^ was used to infer the putative consequences to the structure, splicing, and function of the gene product (transcript and/or peptide) based on synonymous and/or nonsynonymous (ns) changes.

In order to develop genetic markers and build a hypothesis, the significant SNPs which overlapped the major SB-resistance QTL on Chromosome-9 were queried to find association to the observed differentially-expressed candidate genes. These same SNPs were also queried against the 3000 Rice Genome data provided by the Rice SNP Seek project^73, 74^ to profile allelic nature and distribution in the diverse *O. sativa* lines. Some of the SNPs were also queried on the Gramene database against the sequenced genomes of wild species of the *Oryza* genus to infer presence or absence. Finally, peer-reviewed literature was mined to extract, and confirm information on the known leaf sheath blight resistance phenotype for these rice lines.

### Scoring alternative splicing events

SpliceGrapher, a Python-based tool ^75^was used to score alternative splicing events. This tool contains splice-site classifier files for each major reference genome, including *Oryza sativa japonica.* These classifiers are used with the first program (sam_filter.py) in the SpliceGrapher pipeline that intakes aligned experimental files and the provided classifier and outputs alignment files filtered by splice site. Next, the filtered alignment files and the reference gene-annotation file (in the gff file format), is used by the second program (predict_graphs.py) to produce a gff file for each gene and its spliced forms. To compare isoforms detected in our samples to the known reference isoforms, we utilized the program (gene_model_to_splicegraph.py) to generate gffs for each known reference gene. The SpliceGrapher pipeline provided statistics and graphical data files. The types of alternative splicing events generated using SpliceGrapher are alternative 5’ (5’-Alt) and 3’ (3’-Alt) events, exon skipping (ES) and intron retention (IR).

### Population study

The RiceCAP SB2 mapping population was developed as a genetic resource to identify lines containing molecular markers associated with SB resistance^76^. The SB2 population consists of 197 doubled-haploid (DH) lines derived from a cross between the susceptible parent Cocodrie (CCDR)^12^ and the resistant parent MCR010277 (MCR)^13^. The population and the two parents were evaluated for leaf sheath blight disease response on a scale of 0-9, where 0 = no disease and 9 = dead plant. Plants were grown and phenotyped across three years at Crowley, LA and Stuttgart, AR, USA ^26^. The top 10 SB resistant individuals SB2-03 (GSOR200003), SB2-109 (GSOR200109), SB2-134 (GSOR200134), SB2-158 (GSOR200158), SB2-161 (GSOR200161), SB2-174 (GSOR200174), SB2-206 (GSOR200206), SB2-225 (GSOR200225), SB2-259 (GSOR200206), and SB2-272(GSOR200272 and the 10 most susceptible individuals, SB2-13 (GSOR200013), SB2-48 (GSOR200048), SB2-88 (GSOR200088), SB2-99 (GSOR200099), SB2-125 (GSOR200125), SB2-144 (GSOR200144), SB-203 (GSOR200203), SB-255 (GSOR200255), SB-276 (GSOR200276), and SB2-314 (GSOR200314) from the SB2 DH population along with the two parents were screened with 135 candidate nsSNP markers (Table 4, Supplementary Data File 3) identified by Silva et al. 2012 ^22^, including the *OsWAK91* SNP. These markers were located in QTL regions on chromosomes 1, 2, 3, 4, 5, 6, 7, 8, 9, 11 and 12 ^15, 23–29^. An ANOVA analysis was performed to calculate the F values, Bonferroni Hochberg False Discovery Rates (FDR), p values and adjusted R^2^ values.

### DNA extraction, amplification, sequencing and analysis

Total DNA was extracted from seedling tissue using a DNeasy Plant Mini Kit (Qiagen Inc., USA). PCR primers were synthesized by Invitrogen Inc., USA for amplification of the region of interest (Table 5). For PCR amplification, we used Ready PCR Mix, 2x (VWR) following the recommended protocol on a C100 Thermocycler (BioRad). The thermocycler was set for 3 minutes at 95C for initial denaturation, followed by 40 cycles of 30 seconds at 95C for denaturation, 30 seconds at 55C for annealing, 30 seconds at 72C for extension, and 5 minutes at 72C for final extension. Amplified DNA fragments were separated on a 1% agarose gel in 1x TAE buffer. These fragments were extracted from the gel using QIAGEN Gel Extraction and QIAGEN PCR product cleanup kits. Isolated PCR products were Sanger sequenced at the CGRB, OSU. The amplified sequences from the PCR products for each experimental line, including MCR and CCDR, were aligned to the reference Nipponbare genome (IRGSP v1.0) using ClustalW for confirmation of SNPs and to identify and confirm the alleles.

**Table 5.**
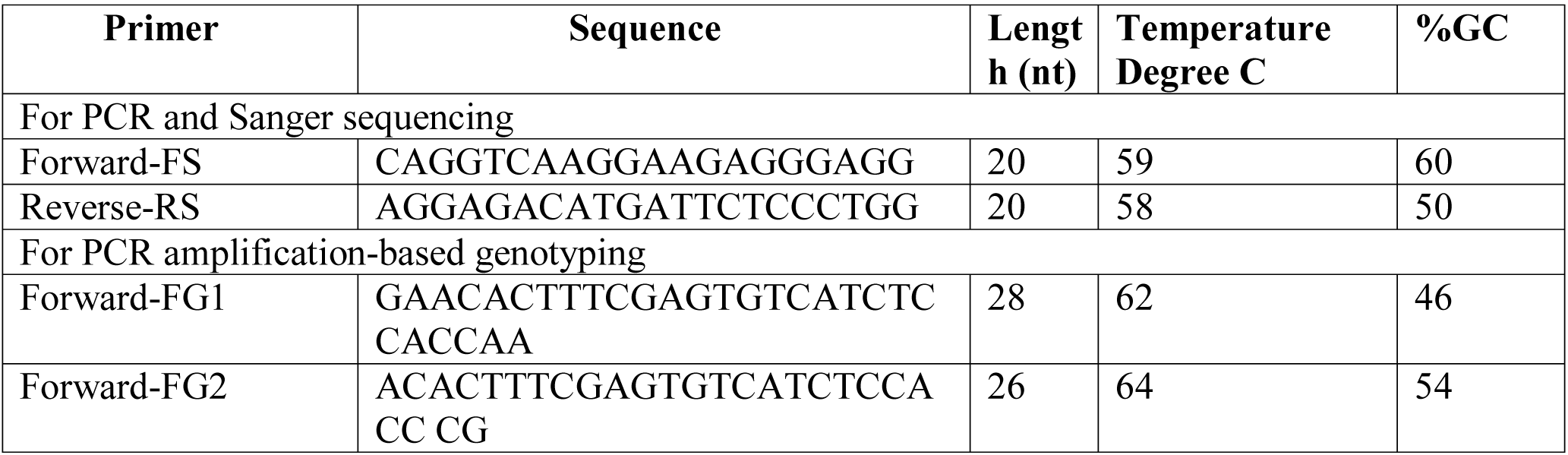
OsWAK SNP Primer Sets and PCR conditions. Primer name with FS and RS suffix were used for amplifying the SNP region and Sanger sequencing. The primer names with suffix FG were used for direct SNP-specific PCR-based genotyping.

### Data mining and functional annotation

The reference genome sequences, gene function and gene family annotations were downloaded in bulk and/or queried online at Gramene database (www.gramene.org) ^44^. The 3000 rice genome project data was mined at Rice SNP-Seek Database (http://snp-seek.irri.org/) ^73, 74^. Gramene database was queried to pull out all rice genes (IRGSP v1.0) annotated to carry the WAK domain and/or were part of WAK gene family. The protein sequences of the canonical (longest) isoform) were downloaded in fasta format to run the CLUSTALW alignment with default parameters. The gene family tree and the differential gene expression data for the respective rice WAK gene family members were uploaded on the iTOL web portal^77^ to generate the graphics (Figure 1K).

## Data Availability

The raw RNA sequence reads were submitted to the EMBL-EBI ArrayExpress (accession number E-MTAB-6402). The raw sequences from the amplified fragments from various rice lines were submitted to the European Nucleotide Archive (study accession PRJEB28811) under the assigned accession numbers ERS2758541-51.

## Acknowledgments

We would like to thank Center for Genome Research and Biocomputing (CGRB) core facility staff, Mark Dasenko for Illumina cluster generation and sequencing, Matthew Peterson, Chris Sullivan and Justin Elser for computational support and Dr. Laurel Cooper and Justin Preece for English language editing. Funding for the project was supported by NSF awards (IOS-1127112 and IOS- 1340112) and the Louisiana Rice Research Board.

## Author Contributions

PJ and JO conceived and designed the experiments. YSG, MG, NAB and AM performed the experiments. NAB, MG, AM, YSG, PJ, JO analyzed the data. Everyone wrote the paper:

## Competing Interest

The Authors declare no competing interests.

