## Supplementary Figure 1 for "Loss of premature stop codon in the *Wall-Associated Kinase 91* (*OsWAK91*) gene confers sheath blight disease resistance in rice"

### Supplementary file-1

#### Chitin Elicitor Binding (OsCEBiP) Gene Family

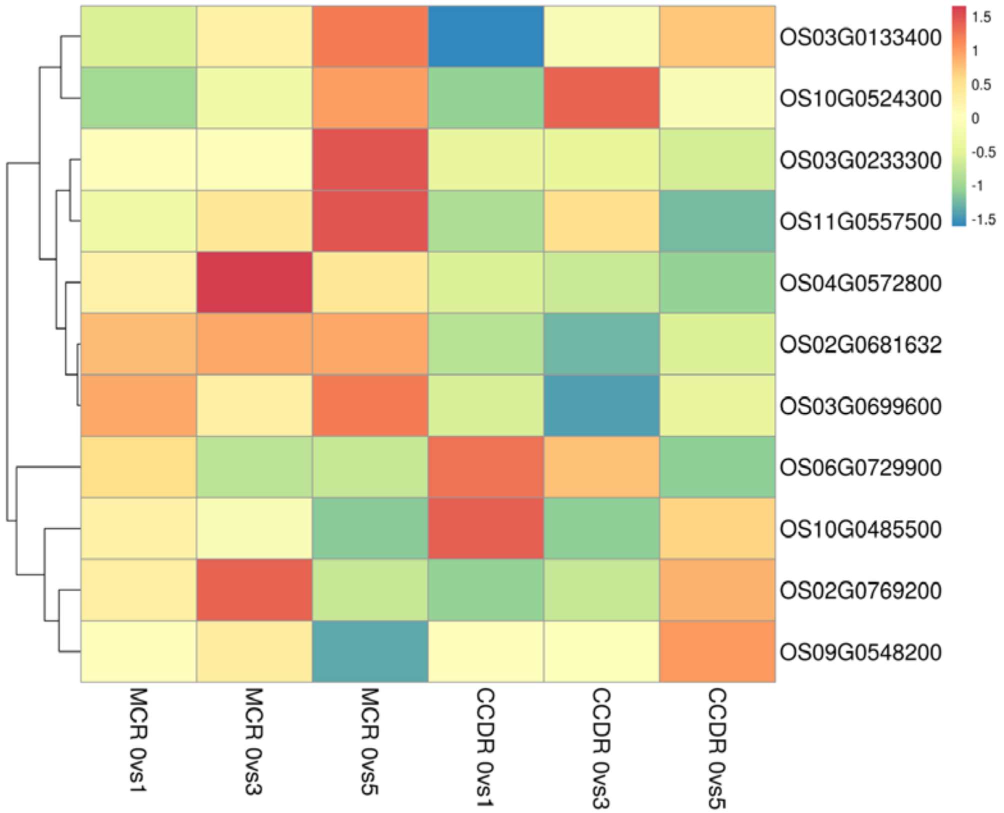


#### Chitin Elicitor Receptor Kinase (CERK) Gene Family

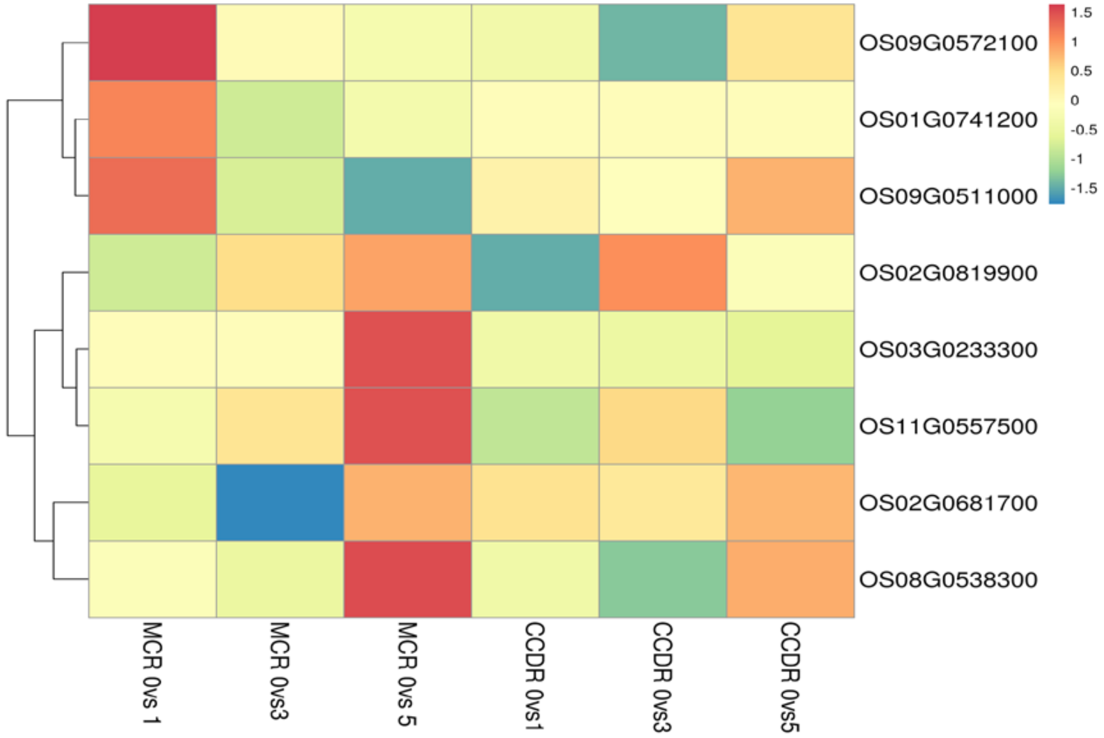


Differential gene expression of the rice Chitin Elicitor Binding (OsCEBiP) and Chitin Elicitor Receptor Kinase (OsCERK) gene family members in the resistant MCR and susceptible CCDR line at Day-1, Day-3 and Day-5 time points after inoculation compared to the Day-0 untreated samples.
